# *Arabidopsis SMN2/HEN2*, Encoding DEAD-box RNA Helicase, Governs Proper Expression of the Resistance Gene *SMN1*/*RPS6* and Is Involved in Dwarf, Autoimmune Phenotypes of *mekk1* and *mpk4* Mutants

**DOI:** 10.1101/2020.05.01.071647

**Authors:** Momoko Takagi, Naoki Iwamoto, Yuta Kubo, Takayuki Morimoto, Hiroki Takagi, Fuminori Takahashi, Takumi Nishiuchi, Keisuke Tanaka, Teruaki Taji, Hironori Kaminaka, Kazuo Shinozaki, Kazuya Akimitsu, Ryohei Terauchi, Ken Shirasu, Kazuya Ichimura

## Abstract

In *Arabidopsis thaliana*, a mitogen-activated protein kinase pathway, MEKK1–MKK1/MKK2–MPK4, is important for basal resistance, and disruption of this pathway results in dwarf, autoimmune phenotypes. To elucidate the complex mechanisms activated by the disruption of this pathway, we have previously developed a mutant screening system based on a dwarf autoimmune line that overexpressed the N-terminal regulatory domain of MEKK1. Here, we report that the second group of mutants, *smn2*, had defects in the *SMN2* gene, encoding a DEAD-box RNA helicase. *SMN2* is identical to *HEN2*, whose function is vital for the nuclear RNA exosome because it provides non-ribosomal RNA specificity for RNA turnover, RNA quality control, and RNA processing. Aberrant *SMN1*/*RPS6* transcripts were detected in *smn2* and *hen2* mutants. Disease resistance against *Pseudomonas syringae* pv. *tomato* DC3000 (*hopA1*), which is conferred by *SMN1*/*RPS6*, was decreased in *smn2* mutants, suggesting a functional connection between *SMN1*/*RPS6* and *SMN2*/*HEN2*. We produced double mutants *mekk1smn2* and *mpk4smn2* to determine whether the *smn2* mutations suppress the dwarf, autoimmune phenotypes of the *mekk1* and *mpk4* mutants, as the *smn1* mutations do. As expected, the *mekk1* and *mpk4* phenotypes were suppressed by the *smn2* mutations. These results suggested that *SMN2* is involved in proper function of *SMN1*/*RPS6*. The GO enrichment analysis using RNA-seq data showed that defense genes were downregulated in *smn2*, suggesting positive contribution of *SMN2* to genome-wide expression of defense-related genes. In conclusion, this study provides novel insight into plant immunity via *SMN2*/*HEN2*, an essential component of the nuclear RNA exosome.

## Introduction

Plant immunity is comprised of pattern-triggered immunity (PTI) and effector-triggered immunity (ETI). Pattern recognition receptors (PRRs) on the cell surface perceive microbe-associated molecular patterns and activate PTI against a broad range of microbes (Dodds and Rathjen, 2010; Ranf, 2017; Tsuda and Katagiri, 2010). Virulent microbial pathogens deploy effectors that suppress PTI and multiply in the plant tissues (Asai and Shirasu, 2015; Dodds and Rathjen, 2010). The recognition of such effectors by nucleotide-binding domain–leucine-rich repeat (NLR) proteins induces ETI, often accompanied by localized cell death, called the hypersensitive response (Cui et al., 2015). The *Arabidopsis thaliana* mitogen-activated protein kinase (MAPK) pathway MEKK1–MKK1/MKK2–MPK4 functions downstream of the PRRs and provides basal resistance against virulent oomycete *Hyaloperonospora arabidopsidis* Noco2 and bacterial pathogen *Pseudomonas syringae* pv. *tomato* DC3000 (Zhang et al., 2012). Disruption of this pathway results in dwarf, autoimmune phenotypes because of constitutive defense responses such as spontaneous cell death and accumulation of reactive oxygen species (Qiu et al., 2008; Suarez-Rodriguez et al., 2010). Study with the *mekk1* mutant showed that the dwarf, autoimmune phenotypes are temperature dependent and partially dependent on *RAR1* or *SID2* (Ichimura et al., 2006). Because NLR activation is often suppressed by high temperature and is suppressed in *rar1* or *sid2* mutants, the loss of *mekk1* has been suggested to activate an NLR pathway(s) (Ichimura et al., 2006). As predicted, screening for genetic suppressors of the dwarf phenotype of *mkk1mkk2* revealed that a coiled-coil-type NLR protein–encoding gene, *SUMM2*, is involved in the dwarf, autoimmune phenotypes caused by the disruption of the MEKK1–MKK1/MKK2–MPK4 pathway (Zhang et al., 2012). The same suppressor screening identified *SUMM1/MEKK2, SUMM3/CRCK3,* and *SUMM4/MKK6* (Kong et al., 2012; Lian et al., 2018; Zhang et al., 2017). Disruption of the MEKK1–MKK1/MKK2–MPK4 pathway removes MPK4-mediated repression of SUMM1/MEKK2, the closest paralog of MEKK1, and constitutively activates SUMM2 (Zhang et al., 2017). MPK4 phosphorylates calmodulin-binding receptor-like cytoplasmic kinase 3 (SUMM3/CRCK3) and a decapping enhancer (PAT1), and these proteins also associate with SUMM2 *in planta* (Roux et al., 2015; Zhang et al., 2017). SUMM3/CRCK3 and PAT1 may serve as a guardee or decoy of SUMM2 (Zhang et al., 2017).

Recently, we generated estradiol-inducible MEKK1N-Myc transformants of Landsberg *erecta* (L*er*) as phenocopies of the *mekk1* mutant. We mutagenized seeds of the transformants and performed genetic screening to isolate suppressors of the MEKK1N-Myc overexpression–induced dwarf phenotype. We chose the 9 mutants with the best dwarfism suppression. The mutant loci were designated *<u>s</u>uppressor of <u>M</u>EKK1<u>N</u> overexpression-induced dwarf* (*SMN*) *1* and *SMN2* (Takagi et al., 2019). We have reported the isolation of five mutants belonging to the first complementation group (Takagi et al., 2019). We have identified a locus, designated as *SMN1*, encoding the TIR-class NLR protein RPS6, which recognizes the HopA1 effector from *Pseudomonas syringae* and is required for resistance to this pathogen (Kim et al., 2009; Takagi et al., 2019). Mutations in *SMN1*/*RPS6* partially suppress the dwarf, autoimmune phenotypes of *mekk1* and *mpk4*. We have suggested that two structurally distinct NLR proteins, SMN1/RPS6 and SUMM2, monitor the integrity of the MEKK1–MKK1/MKK2–MPK4 pathway (Takagi et al., 2019). The mRNA levels of *SMN1*/*RPS6* are regulated by the nonsense-mediated mRNA decay (NMD) surveillance mechanism that degrades aberrant mRNAs in the processing body (P-body) in the cytosol (Gloggnitzer et al., 2014). Impairment of NMD by loss of functional mutation in *SMG7*, encoding NMD component, leads to autoimmunity through deregulation of the *SMN1/RPS6* transcript (Gloggnitzer et al., 2014).

In addition to the P-body in the cytosol, the RNA exosome, located in both the cytosol and nucleus, is a major machinery for RNA degradation. The RNA exosome degrades aberrant RNAs in the 3′ to 5′ direction and is involved in RNA turnover to regulate RNA level, RNA quality control to eliminate defective RNAs, and RNA processing for maturation of precursor RNAs (Chiba and Green, 2009; Jensen, 2010; Zhang and Guo, 2017). The nuclear RNA exosome requires the interaction with RNA helicases for its function (Sikorska et al., 2017); two related DEAD-box RNA helicases, mRNA transport-defective 4 (MTR4) and HUA enhancer 2 (HEN2), associate with the nuclear RNA exosome in *Arabidopsis* (Lange et al., 2014). MTR4 and HEN2 are closely related to each other; MTR4, which localizes mainly in the nucleolus and is required for the degradation of rRNA precursors and rRNA maturation by-products (Lange et al., 2011). In contrast, HEN2, a nucleoplasmic protein, has a distinct function to the MTR4 because HEN2 is dispensable for rRNA maturation. *HEN2* was initially identified in a forward genetic screen for an enhancer of a flower morphology defect in a *hua1hua2* double mutant (Western et al., 2002). Accumulation of a misprocessed *AGAMOUS* transcript in the *hen2* mutant suggests that HEN2 is required for degradation of aberrant *AGAMOUS* transcripts (Cheng et al., 2003). *HEN2* has also been identified as a genetic suppressor (*SOP3*) of a developmental defect caused by a weak *pas2-1* allele encoding 3 hydroxy acyl-CoA dehydratase, which is essential for the very long chain fatty acids elongation (Hématy et al., 2016).

However, very little is known about whether the nuclear RNA exosome regulates innate immunity in plants. In this study, we found that *SMN2*, identical to *HEN2,* is a novel suppressor of the dwarf, autoimmune phenotypes of the *mekk1* and *mpk4* mutants. We suggest that *SMN2* is required for proper function of *SMN1/RPS6*.

## Results

### Identification of the *SMN2* gene by MutMap analysis

To gain insight into the autoimmune and severe dwarf phenotypes of the *mekk1* mutant and to avoid the difficulty of propagating the *mekk1* homozygous mutant, we generated estradiol-inducible MEKK1N-Myc transformants of L*er* as phenocopies of the *mekk1* mutant. Using this transgenic line, we performed genetic suppressor screening to identify gene(s) involved in the autoimmune and severe dwarf phenotypes of the *mekk1* mutant. Here we report three mutants belonging to the second complementation group: *smn2-85*, *smn2-90*, and *smn2-91* (Fig. 1A). To find the causal suppressor gene, we backcrossed the three mutants to the parental line, the estradiol-inducible MEKK1N-Myc transgenic L*er*, and produced F2 individuals from backcross 1 generation with the *smn* phenotype, which were then subjected to a MutMap analysis (Abe et al., 2012). We found that *smn2-90* and *smn2-91* shared almost all of the SNPs. Because these mutants came from identical M2 seed batch, we concluded that *smn2-90* and *smn2-91* are siblings. In the MutMap analysis, the *smn2* mutants shared the highest peak of the SNP index on chromosome 2 (Supplemental Fig. S1). *At2g06990* was the only gene in which nonsynonymous mutations were found in all *smn2* alleles and was thus identified as *SMN2* (Fig. 1B). The *smn2-85* mutant had a G472D amino acid substitution, and both *smn2-90* and *smn2-91* carried a nonsense mutation resulting in a change of W237 to a stop codon (Fig. 1B). We designed CAPS markers for *smn2-85* (Supplemental Table S1) and SNP-specific PCR markers for *smn2-90* (Supplemental Table S2). Using these markers, we removed the estradiol-inducible MEKK1N-Myc transgene by crossing for further analyses.

**Fig. 1.**
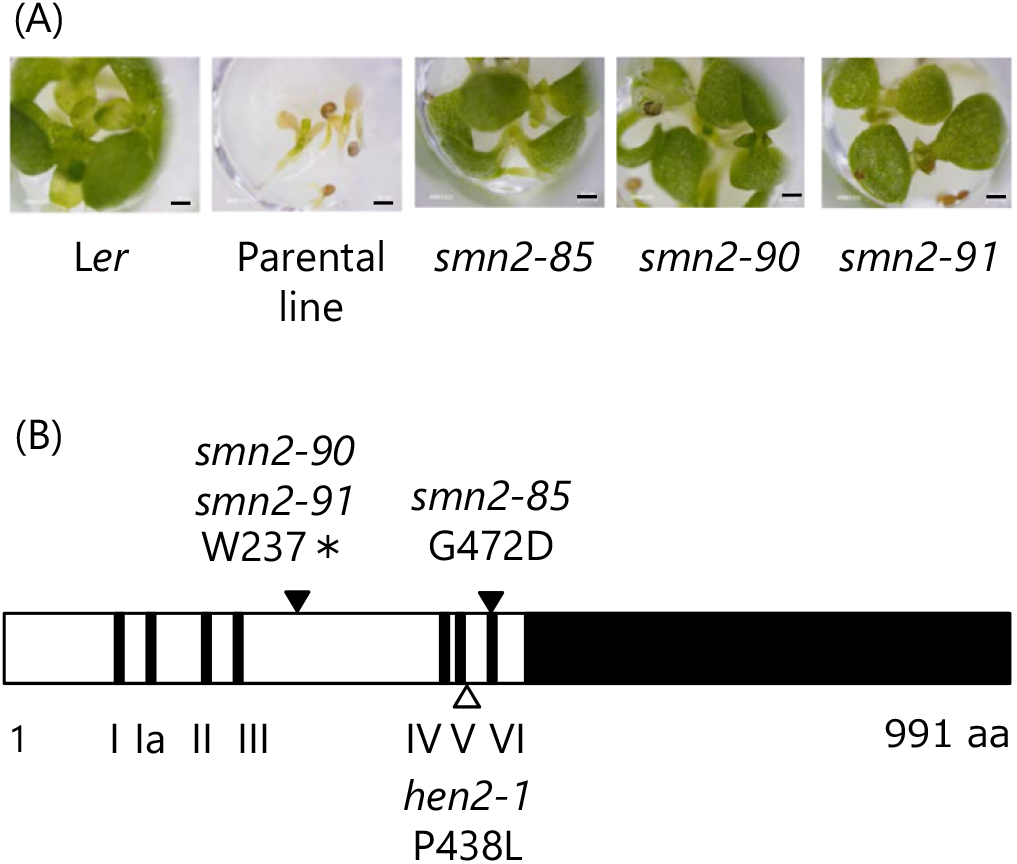
Identification of the gene mutated in the *smn2* mutants. (A) Unlike the parental line, the *smn2* mutants do not show the dwarfism induced by MEKK1N overexpression. Seeds of the wild type (L*er*), the parental line for mutant screening, and *smn2* mutants carrying the estradiol-inducible MEKK1N-Myc transgenes were germinated in liquid GM containing 20 μg/ml estradiol on filter paper for 10 days. Bars = 0.5 mm. (B) Diagram of the SMN2/HEN2 protein. Black bars, helicase motifs I to VI. Triangles, amino acid substitutions encoded by *smn2* alleles (closed) and *hen2-1* (open); individual allele numbers are shown. Asterisk, stop codon. Black box on the right, C-terminal domain.

### Tissue-specific expression of *SMN2*

In the *mekk1* mutants, cell death and H_2_O_2_ accumulation were observed in the vasculature (Ichimura et al., 2006). Therefore, we investigated whether *SMN2* is expressed in the vasculature. A 2-kb DNA fragment containing the *SMN2* promoter was transcriptionally fused to the *GUS* reporter gene and introduced into WT L*er*. The GUS reporter was expressed predominantly in the vascular tissues, especially in expanded cotyledons and true leaves (Fig. 2A, B). Expression of GUS in mesophyll cells was also visible in emerging true leaves and to a lesser extent in expanded leaves (Fig. 2A). This *SMN2* expression pattern overlapped with tissue-specific expression of *MEKK1* (Ichimura et al., 2006). These results suggested a role of *SMN2* in the autoimmune phenotype of *mekk1*. We also found strong GUS expression in the root tip, root stele, and lateral root primordia (Fig. 2C, D).

**Fig. 2.**
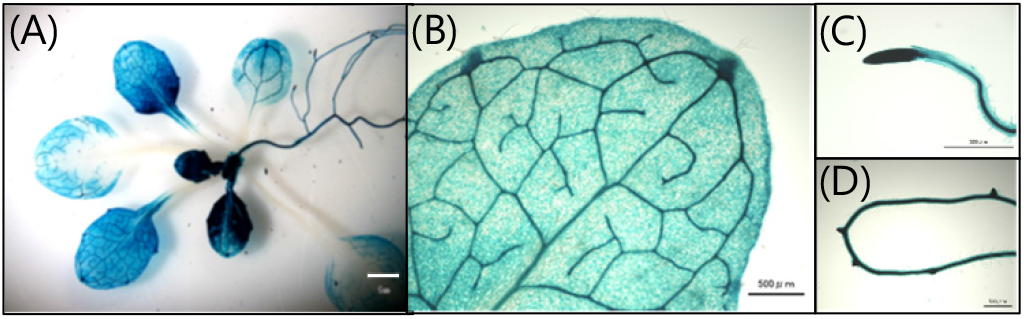
Tissue-specific expression of *SMN2*. A 1.6-kb promoter region of *SMN2* was transcriptionally fused to the GUS reporter gene. The *SMN2* promoter::*GUS* transgenic seeds were germinated and grown on GM medium, and 3-week-old seedlings were stained with X-gluc. One representative out of 12 independent lines is shown. (A) Whole seedling. Bar = 1.0 mm. (B) True leaf. Bar = 500 μm. (C) Root tip. Bar = 500 μm. (D) Lateral root primordia. Bar = 500 μm.

### Aberrant expression of *SMN1/RPS6* in *smn2* mutants

*HEN2* encodes a component of the nuclear RNA exosome targeting complex (Lange et al., 2014). To find whether the expression of any defense genes is altered in the *smn2* mutants, we checked the data obtained by the tilling array analysis of the *hen2* mutant to elucidate target transcripts degraded by the nuclear RNA exosome (Lange et al., 2014). *SMN1/RPS6* was among the 18 defense genes upregulated in the *hen2* mutant (Supplementary Table S3). The *hen2* mutant accumulated an alternative 3′-end transcript and a read-through transcript of *SMN1/RPS6*. We then asked whether aberrant transcripts can be detected in the *smn2* mutants. In the course of Gene Ontology (GO) enrichment analysis (described below), we mapped sequence reads on the *SMN1/RPS6* gene and observed a significant increase in the number of reads at the 3′ UTR and downstream intergenic regions in *smn2-90* in comparison with L*er* (Supplemental Fig. S2). RT-qPCR analysis with primers to amplify the regions corresponding to the probes used in the tilling array analysis (Fig. 3A) confirmed upregulation of *SMN1/RPS6* transcripts at the 3′ UTR and downstream intergenic regions in the *smn2-85* and −90 mutants and *hen2-1*, a mutant originally isolated from L*er* and allelic to *smn2* (Fig. 3B) (Western et al., 2002). Transcript levels in these mutants were almost 5 times that in Ler at site A, 7 times at site B, and >40 times at site C (Fig. 3B). In contrast, the differences between L*er* and the mutants in the levels of *SMN1*/*RPS6* transcripts corresponding to the TIR, NB, and LRR domains were not significant (Supplemental Fig. S3A). This result also suggests possible functional connection between *SMN1/RPS6* and *SMN2/HEN2* via a posttranscriptional mechanism.

**Fig. 3.**
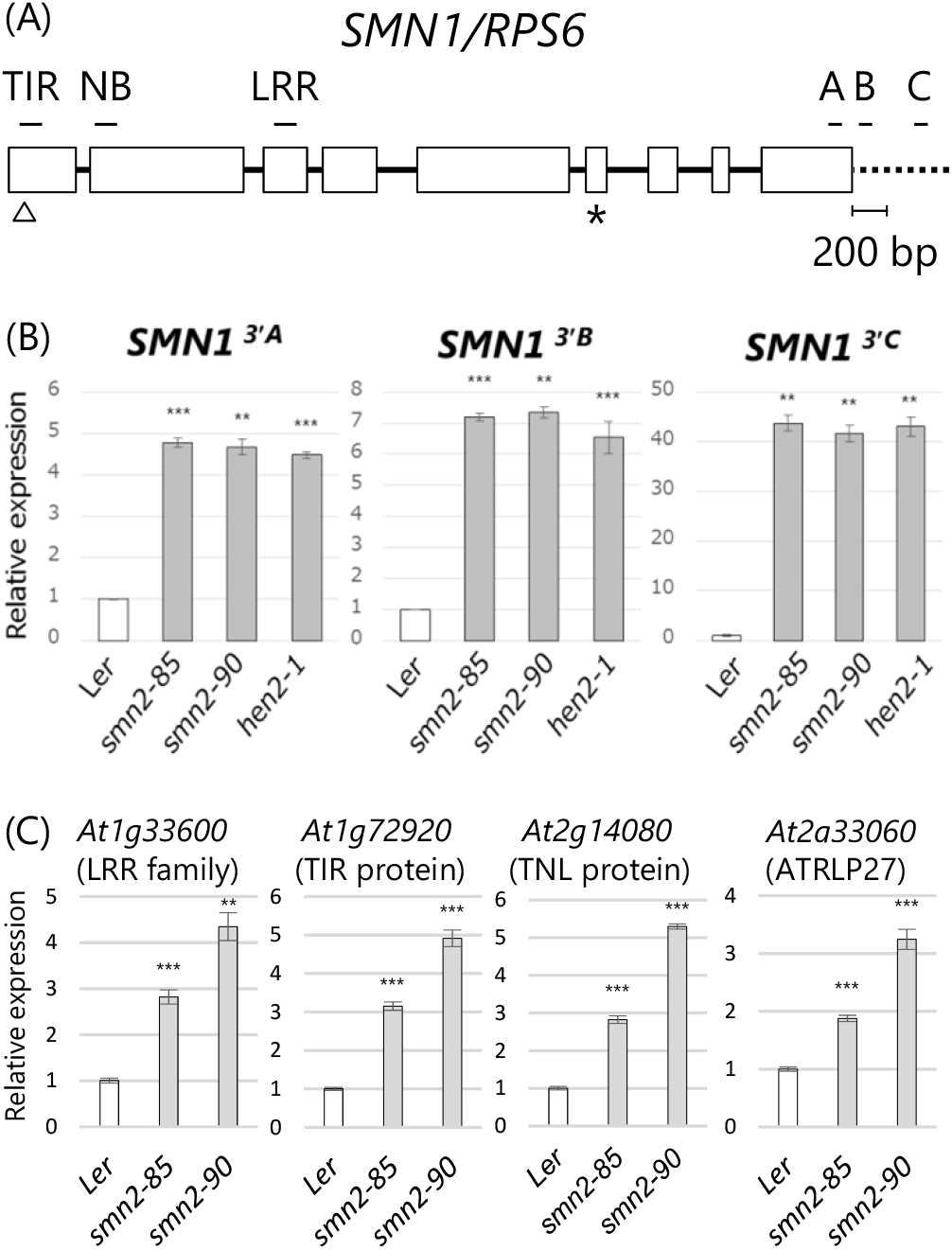
Detection of aberrant transcripts corresponding to 3′ regions of *SMN1/RPS6* and affected defense genes in *smn2* and *hen2-1* mutants. (A) Diagram of the *SMN1*/*RPS6* gene. Open boxes, exons; lines, introns; broken line, 3′ intergenic region; open triangle, initiation codon; asterisk, stop codon. Bars above the TIR, NB, and LRR domains correspond to the RT-qPCR amplicons. Bars under letters “A”, “B”, and “C” correspond to the RT-qPCR amplicons of the 3′ UTR (A) and further downstream intergenic regions B and C. (B, C) Detection of aberrant transcripts corresponding to 3′ UTR and further downstream intergenic regions of (B) *SMN1/RPS6* (regions A to C) and (C) defense genes in the *smn2* and *hen2* mutants. Transcript levels are shown relative to those in L*er* as determined by RT-qPCR using *Actin2* as an internal standard. Plants were grown for 10 days at 22°C. Error bars, SD (n = 3). Representative results from three independent experiments are shown. \*\**p* < 0.005, \*\*\**p* < 0.001 vs. L*er* (Student’s *t* test).

On the basis of the results of the tilling array analysis (Lange et al., 2014; Supplemental Table S3), we selected some other representative defense genes, which encode an LRR family protein (*At1g33600*), TIR protein (*At1g72920*), TNL protein (*At2g14080*), and ATRLP27 (*At2g33060*). Using RT-qPCR, we confirmed accumulation of aberrant transcripts at the 3′-end regions (Fig. 3C), but not the coding regions of these genes, in *smn2* mutants (Supplemental Fig. S3B). We also analyzed transcript levels of the *SUMM2* coding region near the 3′ end because this gene has no 3′ UTR. The *SUMM2* transcript levels in the *smn2* and *hen2* mutants were 1.3 to 1.4 times those in L*er* (Supplemental Fig. S4). Our results confirmed the generation of aberrant transcripts at the 3′ regions of defense genes, including *SMN1/RPS6*, in *smn2* mutants.

### The *smn2* mutations result in loss of function of *SMN1*/*RPS6*

We checked whether abnormal expression of *SMN1/RPS6* in the *smn2* mutants alters resistance conferred by *SMN1/RPS6*. We inoculated the *smn2-85* and *smn2-90* mutants with *Pseudomonas syringae* pv. *tomato* (*Pst*) DC3000 (*hopA1*). We used *smn1-150* as a positive control for the loss of resistance. Bacterial growth was highest in *smn1-150*, and was significantly higher in *smn2-85* and *smn2-90* than in L*er* (Fig. 4), suggesting that *smn2* mutations resulted in loss of function of *SMN1*/*RPS6*. Disease resistance against *Pst* DC3000 was not altered in *smn1* and *smn2* mutants compared to L*er* (Supplemental Fig. S5).

**Fig. 4.**
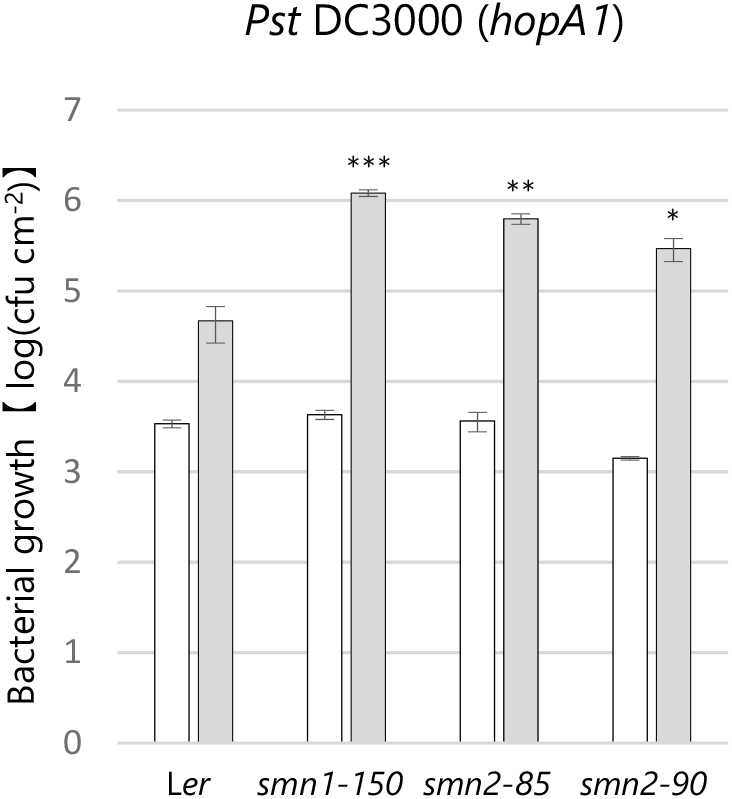
Population growth of *Pst* DC3000 (*hopA1*) in *smn2* mutants. Plants (7 to 8 week old) were inoculated at 2 × 10^4^ cfu/mL. Values are averages for leaf tissue 0 day (white bar) and 3 days (gray bar) after inoculation. Error bars, SD (n = 3). This experiment was performed three times with similar results. \**p* < 0.05, \*\**p* < 0.01, \*\*\**p* < 0.001 vs. L*er* (Student’s *t* test).

### Genome-wide identification of defense genes affected by *smn2* mutations

Using L*er* and *smn2-90*, we performed RNA-seq analysis and detected 250 differentially expressed genes (DEGs). Of those, 86 DEGs were downregulated in *smn2-90*. To elucidate the function of the downregulated genes, we performed GO enrichment analysis with the agriGO. We found that the downregulated genes were involved in response to stimulus (GO:0050896), response to stress (GO:0006950), and defense response (GO:0006952) (Supplemental Fig. S6). To confirm the downregulation of defense genes in *smn2* mutants, we examined four representative genes, *ECS1* (At1G31580), *FMO1* (At1G19250), *WRKY54* (At2G40750), and *WRKY70* (At3G56400), by RT-qPCR with the promiers correspond to the coding regions. The expression levels of *ECS1*, *FMO1*, *WRKY54*, and *WRKY70* in *smn2* mutants were reduced to about 40%, 40% to 56%, 56% to 70%, and 56% in *smn2* mutants, respectively (Fig. 5). These data suggested a positive contribution of *SMN2* to genome-wide expression of defense genes.

**Fig. 5.**
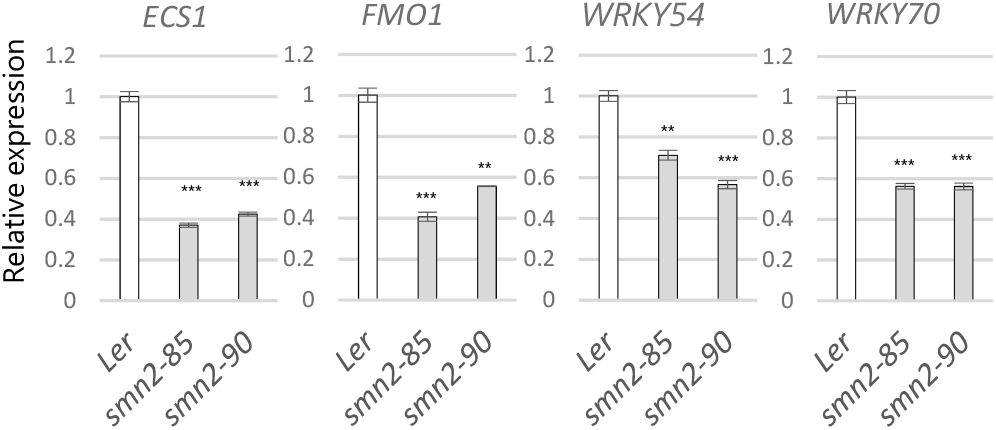
Expression levels of representative defense genes are reduced in *smn2* mutants. Plants were grown for 10 days at 22°C. Transcript levels are shown relative to L*er* as determined by RT-qPCR using *Actin2* as an internal standard. Error bars, SD (n = 3). Representative results from three independent experiments are shown. \*\**p* < 0.01, \*\*\**p* < 0.001 vs. L*er* (Student’s *t* test).

### Suppression of dwarf, autoimmune phenotypes of *mekk1* by *smn2* and *hen2* mutations

To test whether *SMN2* is involved in the dwarf, autoimmune phenotypes of *mekk1*, we generated *mekk1smn2* double mutants. We also used *hen2-1* to increase the accuracy of phenotype analyses. The dwarf phenotype of the *mekk1* mutant was partially suppressed in the *mekk1smn2* and *mekk1hen2* double mutants at 26°C (Fig. 6A). To test whether autoimmune phenotype accompanied by cell death and H_2_O_2_ accumulation in *mekk1* were also suppressed in *mekk1smn2* and *mekk1hen2*, we stained leaf tissues with trypan blue to see cell death or diaminobenzidine (DAB) to see H_2_O_2_ accumulation. Fewer cells were stained with trypan blue or DAB in *mekk1smn2* and *mekk1hen2* than in *mekk1* at 26°C (Fig. 6B, C). In particular, far fewer cells were stained with trypan blue in the mesophyll of *mekk1smn2* and *mekk1hen2* (Fig. 6B). Consistently, DAB staining in the vasculature and adjacent cells of *mekk1* plants was also suppressed by the *smn2* or *hen2* mutations (Fig. 6C). Our data suggest that *SMN2* is involved in the dwarf, autoimmune phenotypes of *mekk1*.

**Fig. 6.**
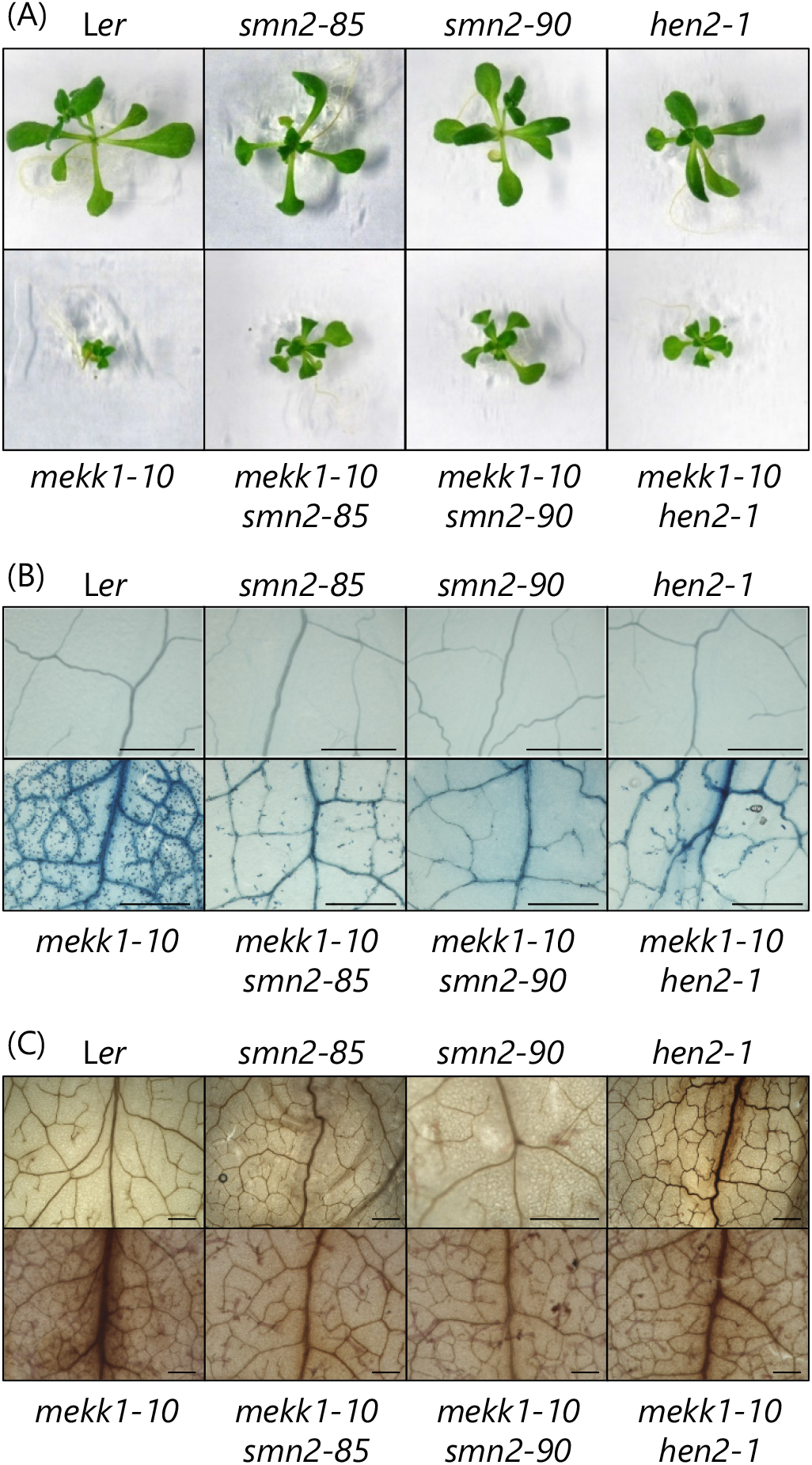
Dwarf, autoimmune phenotypes of the *mekk1* mutant are partially suppressed by the *smn2* and *hen2* mutations. (A) Morphology, (B) cell death, and (C) H_2_O_2_ production. Plants were grown on GM agar for (A) 18 days or (B, C) 14 days at 26°C. Detached true leaves were stained with (B) trypan blue or (C) diaminobenzidine. Bars = 500 μm. Data are representative of at least four biological replicates.

### Suppression of dwarf, autoimmune phenotypes of *mpk4* by *smn2* and *hen2* mutations

*SMN1* is involved in the dwarf, autoimmune phenotype of *mpk4* (Takagi et al., 2019). To examine whether the *smn2* and *hen2* mutations suppress the *mpk4* phenotypes, we produced *mpk4smn2-85* and *mpk4hen2-1* double mutants. The double-mutant plants were clearly larger than the *mpk4* plants, but smaller than L*er*, *smn2*, or *hen2* plants at 24°C (Fig. 7A). Consistently, cell death and H_2_O_2_ accumulation were partially suppressed in the *smn2* and *hen2* mutants in comparison with those of *mpk4* at 22°C (Fig. 7B, C). These results showed that *SMN2* is involved in the dwarf, autoimmune phenotypes of *mpk4*.

**Fig. 7.**
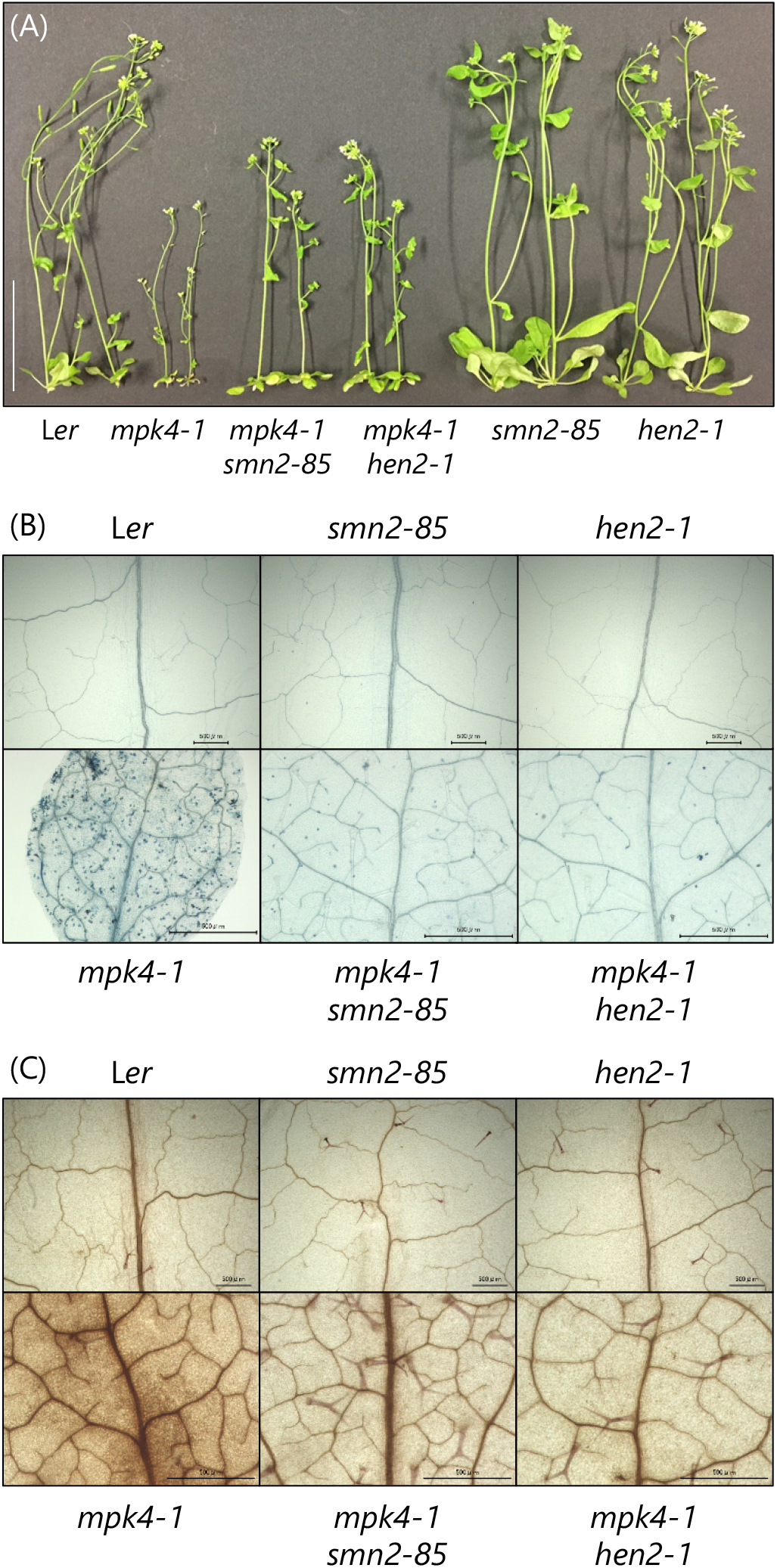
Dwarf, autoimmune phenotypes of the *mpk4* mutant are partially suppressed by the *smn2* and *hen2* mutations. (A) Morphology, (B) cell death, and (C) H_2_O_2_ production. In (A), seeds were germinated on GM for 10 days, and then the plants were grown in soil at 24°C until 1 month of age. Bar = 5 cm. In (B) and (C), plants were grown in soil for 1 month at 22°C. Detached leaves were stained with (B) trypan blue or (C) diaminobenzidine. Bars = 500 μm. Data are representative of three biological replicates.

## Discussion

We have previously reported that loss of function of *SMN1/RPS6*, encoding a TNL immune receptor, partially suppressed the dwarf, autoimmune phenotypes of the *mekk1* and *mpk4* mutants, and suggested that SMN1/RPS6 monitors the integrity of the MEKK1–MKK1/MKK2–MPK4 pathway (Takagi et al., 2019). In this study, we found that *SMN2* is identical to *HEN2* and is the causal gene of the second complementation group of the *smn* mutants. We found that disease resistance conferred by *SMN1*/*RPS6* was decreased by the mutations in *SMN2*/*HEN*. The *smn2* and *hen2* mutations also partially suppressed the dwarf, autoimmune phenotypes of *mekk1* and *mpk4*. We propose that *SMN2* is involved in the dwarf, autoimmune phenotypes caused by disruption of the MEKK1–MKK1/MKK2–MPK4 pathway by regulating *SMN1*/*RPS6* expression.

### Identification of *SMN2/HEN2*

Among the suppressor mutants of the dwarf, autoimmune phenotype induced by conditional overexpression of the N-terminal regulatory domain of MEKK1, we have chosen the nine best-scoring mutants: five *smn1*/*rps6* (Takagi et al., 2019), three *smn2*/*hen2* (this study), and a distinct mutant (to be reported elsewhere). As the *smn2-90* and *smn2-91* mutants originated from the same parent, the *smn2* mutants were actually represented by two alleles, *smn2-85* and *smn2-90*. Fewer *smn2* alleles compared to five *smn1* alleles may correspond to the degree of contribution of wild-type *SMN1* and *SMN2* to the dwarf phenotype of MEKK1N-Myc-expressing plants. Sequence analysis showed that *smn2-85* had a G472D amino acid substitution, and that both *smn2-90* and *smn2-91* carried a nonsense mutation resulting in a stop codon instead of W237. The residue mutated in *smn2-85* is the first of the two glycines in the highly conserved motif VI (Y_467_IQMS<u>G</u>RAGR_476_), required for RNA binding and ATP hydrolysis (Aubourg et al., 1999; Pause et al., 1993). An in vitro binding assay of the yeast initiation factor 4A (eIF-4A), a DEAD-box RNA helicase, suggested that R362 in motif VI is essential for interaction to both RNA and ATP (Pause et al., 1993). The R362 residue of the yeast eIF-4A correspond to R473 in motif VI of SMN2/HEN2. The G472D substitution in *smn2-85* adjacent to this critical amino acid could alter charge distribution in motif VI, possibly hampering the interaction with both RNA and ATP. The nonsense mutation in *smn2-90* and *smn2-91* between motifs III and IV is also very likely to cause loss of function of SMN2/HEN2.

### Possible connection between nuclear RNA exosome and defense responses

The nuclear RNA exosome contains a nine-subunit core (Exo9). Six subunits are similar to RNase PH, but the Exo9 in yeast and humans has lost the active sites, and Rrp44 is tightly associated with Exo9 to degrade RNA (Jensen, 2010). *Arabidopsis* Exo9 has RNase activity but requires the interaction with exosome-associated DEAD-box RNA helicases for its function (Sikorska et al., 2017). In *Arabidopsis*, two related DEAD-box RNA helicases, MTR4 and HEN2, associate with the nuclear RNA exosome (Lange et al., 2011). In contrast to MTR4, HEN2 is important for degradation of non-ribosomal RNAs (Lange et al., 2014). Loss of function of *HEN2* results in accumulation of polyadenylated nuclear RNAs (exosome substrates) such as short transcripts derived from mRNA genes, 3′-extended mRNAs, incompletely spliced mRNAs, snoRNA and miRNA precursors, and spurious transcripts produced from pseudogenes and intergenic regions (Lange et al., 2014).

In this work, we detected aberrant transcripts of the 3′ UTR of *SMN1/RPS6* in the *smn2-90* mutant (Supplemental Fig. S2), but no differences in the levels of coding region transcripts (Supplemental Fig. S3A). We also validated the occurrence of aberrant RNA derived from the 3′ regions of 4 genes (Fig. 3C) selected from among 18 defense genes (Supplemental Table S3) detected in the tilling array analysis with *hen2* by Lange et al. (2014). Our GO enrichment analysis with the RNA-seq data showed that the majority of genes downregulated in *smn2-90* were stress- and/or defense-related, suggesting that *SMN2*/*HEN2* has broad role(s) in mRNA degradation or RNA processing in response to such stimuli. However, inoculation of the *smn2* mutants and L*er* with *Pst* DC3000 resulted in similar disease resistance (Supplemental Fig. S5). The effect of *smn2* mutations on defense gene expression may be below the threshold to alter disease resistance, or may become detectable with other pathogens. Our findings unveil a possible function of SMN2/HEN2 in posttranscriptional gene regulation in biotic stress responses in plants.

The barley *RRP46* gene, encoding a component of the RNA exosome, is responsible for the phenotype of the *Blumeria graminis* f. sp. *hordei (Bgh)-induced tip cell death1* (*bcd1*) mutant (Xi et al., 2009). Deletion or knockdown of this gene results in infection-induced tip cell death regardless of *Bgh* isolate, suggesting a potential link between RNA exosome and defense responses. Microarray analysis revealed that the function of most genes overexpressed in *bcd1* is associated with ribosomal RNAs and proteins (Xi et al., 2009). *Arabidopsis* mutant phenotypes of *RRP46* (At3G46210) and its paralog (At2g07110) have not been reported.

### 3′ region–derived aberrant transcripts normally degraded by nuclear RNA exosome abolish the function of *SMN1*/*RPS6*

The *smn2* and *hen2* mutations led to a production of abnormal mRNAs relevant to the phenotypes of the *agamous* (Cheng et al., 2003) and *pas2-1* mutants (Hématy et al., 2016). The *hen2-1* mutant accumulates aberrant *AGAMOUS* transcripts containing at least the first intron and a large portion of the second intron, suggesting that *HEN2* is involved in the degradation of misprocessed *AGAMOUS* transcripts (Cheng et al., 2003). The *pas2-1* mutation results in *pas2-1* mRNA variants that are degraded by the nuclear RNA exosome. In the double *pas2-1 suppressor of pas2-1* (*sop*) *3* mutant, one of the *pas2-1* mRNA isoforms producing functional PAS2 protein was stabilized, resulting in restoration of the developmental defect; the gene mutated in *sop3* is identical to *HEN2* (Hématy et al., 2016). The *smn2* and *hen2* mutations resulted in production of *SMN1/RPS6* transcripts with shortened or extended 3′ UTRs, which are normally degraded by the nuclear RNA exosome (this study and Lange et al., 2014), resulting in loss of function. This phenomenon is distinct from the cases of *AGAMOUS* and *pas2-1*. The exact step between the transcription and translation of *SMN1/RPS6* affected by the aberrant *SMN1/RPS6* RNAs needs to be elucidated. These data strongly suggest the importance of the nuclear RNA exosome in proper gene expression in plants.

The 3′ UTRs of mRNAs regulate their localization, translation, and stability. Dehydration stress extends the mRNA 3′ UTRs of a subset of stress-related genes, and the extended 3′ UTRs act as repressors or activators, or lead to read-through transcription (Sun et al., 2017). An RNA-binding polyadenylation regulatory factor, FPA, contributes to the extension of 3′ UTR. FPA negatively regulates flg22-induced reactive oxygen species burst and disease resistance against *Pst* DC3000 by producing different isoforms of the defense-related transcriptional repressor ERF4 through choosing alternative polyadenylation sites (Lyons et al., 2013). The transcript level of *ERF4* is increased in *hen2-1* (Lange et al., 2014). Thus, the stress-induced extension of mRNA 3′ UTRs may happen not only in response to dehydration, and SMN2/HEN2 might contribute to these phenomena and finely tune stress-regulated gene expression.

### Suppression of dwarf, autoimmune phenotypes of *mekk1* and *mpk4* by *smn2* and *hen2* mutations

We have previously suggested that SMN1/RPS6 is required for monitoring the MEKK1–MKK1/MKK2–MPK4 pathway (Takagi et al., 2019). Disruption of this pathway by pathogen effector(s) or by T-DNA or transposon insertion activates SMN1/RPS6 and SUMM2 and induces dwarfism and constitutive defense responses (Takagi et al., 2019). In this study, we showed that the *smn2* and *hen2* mutations resulted in loss of function of *SMN1*/*RPS6* with occurrence of aberrant transcripts at the 3′ region of *SMN1*/*RPS6*. These mutations suppressed the dwarf, autoimmune phenotypes of the *mekk1* and *mpk4* mutants. We assume that *smn2*-dependent suppression of the autoimmune phenotypes caused by disruption of the MEKK1–MKK1/MKK2–MPK4 pathway was mediated through the loss of function of *SMN1*/*RPS6*. This suggests that *SMN2/HEN2* plays a novel role in innate immunity by regulating proper *SMN1*/*RPS6* gene expression.

In conclusion, we showed that the plant-specific DEAD-box RNA helicase SMN2/HEN2 is involved in the dwarf, autoimmune phenotypes of *mekk1* and *mpk4*. *SMN2*/*HEN2* is also required for disease resistance mediated by *SMN1*/*RPS6*. We here revealed novel lines of evidence for the involvement of the nuclear RNA exosome in plant immunity via SMN2/HEN2. A genome-wide approach to isolate RNAs bound to SMN2/HEN2 and to elucidate the dynamics of the SMN2/HEN2-bound RNA population before and after pathogen inoculation would allow us to better understand the function of SMN2/HEN2 in innate immunity. This strategy would also be crucial for unraveling novel aspects of innate immunity from the viewpoint of posttranscriptional gene regulation in plants.

## Materials and Methods

### Plant materials, growth conditions, mutant isolation, and MutMap analysis

*Arabidopsis thaliana* plants were grown at 22°C under a 16-h light/ 8-h dark photoperiod in soil or on solid GM medium supplemented with 1% sucrose unless otherwise stated. We used the Landsberg *erecta* (L*er*) accession as WT plants. The mutants in the L*er* background were kindly provided by H. Vaucheret (*hen2-1*), Cold Spring Harbor Laboratory (*mekk1-10*; CSHL DsGT_19053), and H. Hirt (*mpk4-1*; CS5205). To generate the double mutants *mekk1-10smn2-85*, *mekk1-10smn2-90*, and *mekk1-10hen2-1*, *mekk1-10* was crossed with *smn2-85*, *smn2-90*, or *hen2-1*, respectively. To generate the double mutants *mpk4-1smn2-85* and *mpk4-1hen2-1*, *mpk4-1* was crossed with *smn2-85* or *hen2-1*, respectively. Mutant isolation and MutMap analysis are described in Takagi et al. (2019).

### Construction of transgenic plants carrying the *SMN2*/*HEN2* promoter::GUS construct

A genomic DNA fragment of *SMN2*/*HEN2* containing 1634 bp of the 5′ region from the initiation codon was amplified by PCR and cloned into pBI101, which carries the *GUS* coding sequence but no promoter, using the *Hin*dIII and *Xba*I sites with the In-Fusion HD Cloning Kit (Takara Bio, Otsu, Shiga, Japan). The construct was verified by sequencing and used to transform L*er* by the floral dip method (Clough and Bent, 1998). Transformants were selected on GM medium containing 50 μg/mL kanamycin and subjected to histochemical GUS assay.

### Pathogen and plant inoculation

*Pseudomonas syringae* pv. *tomato* strain (*Pst*) DC3000 carrying the empty vector pML123 and *Pst* DC3000 (*hopA1*) carrying the plasmid pLN92 expressing *hopA1* from *P. syringae* strain 61 (Gassmann, 2005) were kindly provided by W. Gassmann. Plants were grown in soil at 22°C under an 8-h light/16-h dark photoperiod for 6 to 8 weeks. Rosette leaves were infiltrated with a bacterial suspension of 2 × 10^4^ colony-forming units (cfu)/ml (*Pst* DC3000 (*hopA1*)) or 1 × 10^5^ cfu/ml (*Pst* DC3000) in sterilized distilled water (sDW) using a needleless syringe. Leaf discs (ⵁ 4 mm) were excised on days 0 and 3 after inoculation and crushed in sDW. The homogenates were serially diluted with sDW, plated on selective medium, and incubated at 28°C for 2 days.

### Staining tissues with X-gluc, trypan blue, and DAB

Plants were grown at 22°C for 21 days or 26°C for 14 days on GM medium. Leaves or whole plants were stained with X-gluc for the GUS assay, trypan blue, or DAB (all reagents from Sigma-Aldrich, St. Louis, MO, USA) and observed as in Ichimura et al. (2006).

### RT-qPCR analysis

Plants were grown for 14 days on GM medium at 22°C. Total RNA was isolated from whole plants with a Sepazol RNA I Super G reagent (Nacalai Tesque, Kyoto, Japan) according to the manufacturer’s instructions. First-strand cDNA was synthesized using ReverTra Ace qPCR RT MasterMix with gDNA Remover (Toyobo, Osaka, Japan). RT-qPCR was performed using a StepOnePlus Real-Time PCR system (Applied Biosystems, Foster City, CA, USA) or a CFX Connect Real-Time System (Bio-Rad Laboratories, Hercules, CA, USA) with primers listed in Supplemental Table S4. *Arabidopsis Actin2* was used as an internal control. Relative expression values were calculated using the ddCt method.

### RNA-seq analysis

Two biological replicates of the wild-type plants and the *smn2-90* mutant were used for cDNA library construction and sequencing. Plants were grown for 14 days on GM medium at 22°C. Total RNA was isolated from whole plants with a NucleoSpin RNA Plant Kit (Takara Bio) according to the manufacturer’s instructions. A Dynabeads mRNA Purification Kit was used to purify mRNA and an Ion Total RNA-Seq Kit v2 was used to construct cDNA libraries (both kits from Thermo Fisher Scientific, Beverly, MA, USA). The quality of total RNA, mRNA, and cDNA libraries was analyzed with an Agilent TapeStation (Agilent Technologies, Santa Clara, CA, USA). The cDNA libraries were pooled for emulsion PCR using an Ion PI Hi-Q Chef Kit (Thermo Fisher Scientific). The enriched samples were loaded onto an Ion PI chip v3 with Ion Chef, and sequenced with an Ion Proton instrument (Thermo Fisher Scientific). The raw reads from the libraries were filtered to remove adaptor sequences and low-quality bases, and the clean reads from each library were mapped to the reference *Arabidopsis* genome (TAIR 10) using CLC Genomics Workbench (version 9.0; CLC Bio, Aarhus, Denmark). Expression values were measured as RPKM (reads per kilobase of the transcript per million mapped reads). Statistical significance of the differences in gene expression was assessed by comparing the RPKM values using empirical analysis of digital gene expression in R (edgeR) test (Robinson et al., 2010). The raw sequence reads were deposited in the National Center for Biotechnology Information (http://www.ncbi.nlm.nih.gov/) under the accession number GSE143520.GO enrichment analysis was performed by the agriGO (version 2.0) software (http://systemsbiology.cau.edu.cn/agriGOv2/) (Tian et al., 2017).

## Acknowledgements

We thank Alexander Graf, David Greenshields, Catarina Casais, Kaori Takizawa, and Gang-Su Hyon for technical assistance; Dr. Walter Gassmann for providing *Pseudomonas* strains; Dr. Hervé Vaucheret for providing *hen2-1* seeds; Dr. Keiichi Mochida for data analysis; and Drs. Mitsuru Akita and Susumu Mochizuki for discussion. We thank Prof. Nam-Hai Chua of Rockefeller University, New York, for providing the pER8 vector. We also acknowledge the technical expertise of the Gene Research Center facility, Kagawa University.

## Supplementary Data

**Supplemental Figure S1. SNP index plots of chromosome 2 from the MutMap analysis of *smn2* mutant alleles.**

Green points, individual SNPs. Red lines, sliding-window average of the ΔSNP index (window size, 2 Mb; slide size, 200 kb).

**Supplemental Figure S2. Accumulation of aberrant transcripts of the 3′ region of *SMN1* in *smn2-90* and L*er*.**

Screenshots of RNA-seq data mapped to the *SMN1*/*RPS6* region of the reference *Arabidopsis* genome (TAIR 10) visualized in the Integrative Genomics Viewer (version 2.7.2). Two biological replicates of each L*er* and *smn2-90* were used for RNA-seq. The gene models of *SMN1*/*RPS6* (top) and its coding region (bottom) are indicated in indigo blue, where boxes are exons and lines are introns. Stacked gray objects, reads; stacked sky-blue objects, gaps. Purple dots, deletions; dots of other colors, substitutions vs. reference genome.

**Supplemental Figure S3. Levels of transcripts of the coding regions of *SMN1/RPS6* and affected defense genes in *smn2* and *hen2-1* mutants.**

(A) Transcripts encoding the TIR, NB, and LRR domains of SMN1/RPS6 (see Fig. 3A).

(B) Transcripts of the coding regions of affected defense genes. RNA was extracted from 10- or 14-day-old seedlings, and the transcript levels were analyzed by RT-qPCR. Transcript levels are shown relative to those in L*er*; *Actin2* was used as an internal standard. Error bars, SD (n = 3). A representative result from three independent experiments is shown.

**Supplemental Figure S4. *SUMM2* transcript levels in *smn2* and *hen2-1* mutants.**

Transcript levels of the *SUMM2* coding region near the 3′ end were analyzed by RT-qPCR using RNA extracted from 10-day-old seedlings. Transcript levels are shown relative to those in L*er*; *Actin2* was used as an internal standard. Error bars, SD (n = 3). A representative result from three independent experiments is shown.

**Supplemental Figure S5. Population growth of *Pst* DC3000 in *smn2* mutants.**

Plants (7 to 8 week old) were inoculated at 1 × 10^5^ cfu/mL. Values are averages for leaf tissue 0 day (white bar) and 3 days (gray bar) after inoculation. Error bars, SD (n =3). This experiment was performed three times with similar results.

**Supplemental Figure S6. Gene Ontology (GO) enrichment analysis based on RNA-seq**.

GO term enrichment in the biological process category was analyzed with the agriGO (version 2.0) software (http://systemsbiology.cau.edu.cn/agriGOv2/). Boxes show the GO terms, adjusted *P*-values, GO descriptions, item number mapping the GO in the query list and background, and total number of query list and background. We ranked 28,362 genes according to their differential expression and used them as input for the analysis. The resulting enriched GO terms are visualized using a directed acyclic graph. The degree of color saturation of each node is positively correlated with the significance of the enrichment of the GO term. Non-significant GO terms are shown as white boxes. Branches of the GO hierarchical tree containing no significantly enriched GO terms are not shown. Arrows show connections between different GO terms (see figure key for arrow types).

