## Supplemental Tables S1 - S4 for "*Arabidopsis SMN2/HEN2*, Encoding DEAD-box RNA Helicase, Governs Proper Expression of the Resistance Gene *SMN1*/*RPS6* and Is Involved in Dwarf, Autoimmune Phenotypes of *mekk1* and *mpk4* Mutants"

Supplemental Table S1. CAPS markers for *smn2-85* and *hen2-1*.

| Mutant allele | Primer name | Sequence | Restriction enzyme | Product size |
|---|---|---|---|---|
| *smn2-85* | ONI7_CAPS_smn85 (FP) | 5'- CAG ATG AAG AGA AAG AAG TTG TGG AAC AGG TG -3' | *BsrB*I | WT: 1112/512 bp |
| | ONI8_CAPS_smn85 (RP) | 5'- CGA TCT GGG TAC AAC CTA TGA CCA TAG CTA GAG AGG -3' | | Mutant: 1624 bp |
| *hen2-1* | ONI7_CAPS_smn85 (FP) | 5'- CAG ATG AAG AGA AAG AAG TTG TGG AAC AGG TG -3' | *Aci*I | WT: 718/393/332/180 bp |
| | ONI8_CAPS_smn85 (RP) | 5'- CGA TCT GGG TAC AAC CTA TGA CCA TAG CTA GAG AGG -3' | | Mutant: 718/512/393 bp |

Supplemental Table S2. SNP-specific PCR markers for *smn2-90* and *smn2-91*.

| WT/Mutant allele | Primer name | Sequence |
|---|---|---|
| WT | ONI13 smn9091 co-do WT (FP) | 5'- GCT ACT GAA TTT GCG GAA TGG -3' |
| | ONI4 smn2-90 seq (RP) | 5'- GAT CGG CGC CAC AAG CAG CAA CAT AGG AC -3' |
| *smn2-90*, *smn2-91* | ONI14 smn9091 co-do Mut (FP) | 5'- GCT ACT GAA TTT GCG GAA TGA -3' |
| | ONI4 smn2-90 seq (RP) | 5'- GAT CGG CGC CAC AAG CAG CAA CAT AGG AC -3' |

Supplemental Table S3. Aberrant transcripts of defense-related genes in the hen2 mutant (excerpt from Supplemental Table S4 in Lange et al., 2014). “3' end”, “5' end”, and “central” mean detection of upregulated short transcripts. RT means read-through transcript.

| AGI code | Annotation | Altered transcripts |
|---|---|---|
| AT1G26150 | ATPERK10 | 5' end |
| AT1G19394 | Wall-associated kinase family | overlapping genes |
| AT1G33600 | Leucine-rich repeat (LRR) family protein | alternative 3' end or RT |
| AT1G58602 | LRR and NB-ARC domains-containing disease resistance protein | mis-spliced or introns 1 and 2 |
| AT1G72910 | Toll-Interleukin-Resistance (TIR) domain-containing protein | 3' end |
| AT1G72920 | Toll-Interleukin-Resistance (TIR) domain-containing protein | 3' end |
| AT1G72940 | Toll-Interleukin-Resistance (TIR) domain-containing protein | 3' end |
| AT2G30750 | CYP71A12 | 3' end |
| AT2G14080 | Disease resistance protein (TIR-NBS-LRR class) family | alternative 3' end or RT |
| AT2G15080 | receptor like protein 19 | alternative 5' end or 5' extended mRNA |
| AT2G33060 | ATRLP27, RECEPTOR LIKE PROTEIN 27 | 3' end |
| AT3G16720 | ATL2 | central |
| AT4G01250 | WRKY22 | 3' end |
| AT4G31354 | Necrosis- and ethylene-inducing peptide | comprising both mRNAs and RT |
| AT5G26920 | CBP60G | 3' end |
| AT5G46470 | RPS6/SMN1 TIR-NBS-LRR class resistance protein | alternative 3' end or RT |
| AT5G47230 | ERF5 | 3' end |
| AT5G57220 | CYP81F2 | central |

Supplemental Table S4. Primer information

| Primer name | Target | Sequence |
|---|---|---|
| ONI7_CAPS_smn85 (FP) | *smn2-85, hen2-1* | 5'- CAG ATG AAG AGA AAG AAG TTG TGG AAC AGG TG -3' |
| ONI8_CAPS_smn85 (RP) | *smn2-85, hen2-1* | 5'- CGA TCT GGG TAC AAC CTA TGA CCA TAG CTA GAG AGG -3' |
| ONI13 smn9091 co-do WT (FP) | *smn2-90* | 5'- GCT ACT GAA TTT GCG GAA TGG -3' |
| ONI14 smn9091 co-do Mut (FP) | *smn2-90* | 5'- GCT ACT GAA TTT GCG GAA TGA -3' |
| ONI4 smn2-90 seq (RP) | *smn2-90* | 5'- GAT CGG CGC CAC AAG CAG CAA CAT AGG AC -3' |
| OKI446 (FP) | *ACT2* | 5'- GGT AAC ATT GTG CTC AGT GGT GG -3' |
| OKI447 (RP) | *ACT2* | 5'- AAC GAC CTT AAT CTT CAT GCT GC -3' |
| OMN34 (FP) | *SMN1* (3'A) | 5'- GAG GCA TGT GTA TAT CAG TGG AG -3' |
| OMN35 (RP) | *SMN1* (3'A) | 5'- AAT TGT ACC AGT TCG GAG AAT GA -3' |
| OMN36 (FP) | *SMN1* (3'B) | 5'- CTG AAT TTG TTG GCA TCG AAG A -3' |
| OMN37 (RP) | *SMN1* (3'B) | 5'- CCT ATC TTG AAC TTA GTC ATC GTT TAT C -3' |
| OMN38 (FP) | *SMN1* (3'C) | 5'- GAT AAA TTG GAA GCT TAC GCA GAA -3' |
| OMN39 (RP) | *SMN1* (3'C) | 5'- TGT TCA ACT TCC TTC GTC TCT C -3' |
| OMT146 (FP) | $SMN1^{TIR}$ | 5'- CCA GAA TCT CCT CTC TTT ATG GC -3' |
| OMT147 (RP) | $SMN1^{TIR}$ | 5'- GAT CAG TTT CCT ATC CAG TTC -3' |
| OMT148 (FP) | $SMN1^{NB}$ | 5'- GAT AAC GAA GCA AGC ATG ATC G -3' |
| OMT149 (RP) | $SMN1^{NB}$ | 5'- TCC CGA CCA TCC TCA CTT CCT C -3' |
| OMT150 (FP) | $SMN1^{LRR}$ | 5'- GTT AGA TGG CAC TTA CCG -3' |
| OMT151 (RP) | $SMN1^{LRR}$ | 5'- GAC ATT TCG ATG GGT ATC TG -3' |
| OMT1t (FP) | *SUMM2* (3') | 5'- CTC AGA AGA GTT GAA ACA TCG AAT ATC -3' |
| OMT2t (RP) | *SUMM2* (3') | 5'- GAA ATC AGC CGG TCA CAA ATC -3' |
| OMT63ti (FP) | *AT1G33600* (3') | 5'- CCT AAC ACT GAG CAG CAC TT -3' |
| OMT64ti (RP) | *AT1G33600* (3') | 5'- CTG TTA GTT AGT GTT CTG TTC TGT TAT C -3' |
| OMT65ti (FP) | *AT1G72920* (3') | 5'- TGA GTA GCC AGC CAA TCA ATA A -3' |
| OMT66ti (RP) | *AT1G72920* (3') | 5'- CAC AGC TAA CTA GTG GTG TCA A -3' |
| OMT69ti (FP) | *AT2G14080* (3') | 5'- GAT CGA CAC TAT GAA CGT CTG G -3' |
| OMT70ti (RP) | *AT2G14080* (3') | 5'- GAT TGG CTC TTC TTT GCT CTT TG -3' |
| OMT71ti (FP) | *ATRLP27* (3') | 5'- CTC TGG ATT CAA GAT GGG ATA G -3' |
| OMT72ti (RP) | *ATRLP27* (3') | 5'- AAA CCT CTT TCT TAC ATC AGG T -3' |
| OMT139ti (FP) | *AT1G33600* (CDS) | 5'- GTT CTT CAA CCT AGC GCA TAA C -3' |
| OMT140ti (RP) | *AT1G33600* (CDS) | 5'- GAT CGA GGC TTT CTA GTC TCT C -3' |
| OMT141ti (FP) | *AT1G72920* (CDS) | 5'- GGC GTG GAT CCT TGT CAT T -3' |

| | | |
|---|---|---|
| OMT142ti (RP) | *AT1G72920* (CDS) | 5'- TAC TTT CTC ATG ATC TTC TTC TCT AG -3' |
| OMT145ti (FP) | *AT2G14080* (CDS) | 5'- GAT ACC GAA GCT GCC ATG AT -3' |
| OMT146ti (RP) | *AT2G14080* (CDS) | 5'- AGC TCC CAT CCC AAC TAA AC -3' |
| OMT147ti (FP) | *ATRLP27* (CDS) | 5'- CAT ACT GTC GGT GCT CTT ACT G -3' |
| OMT148ti (RP) | *ATRLP27* (CDS) | 5'- ACT GTG TCA GTG CTT GAA TCT -3' |
| OMT85go (FP) | *PDF1.2c* | 5'- AAA CGC CTG ACC ATG TCC -3' |
| OMT86go (RP) | *PDF1.2c* | 5'- GCT ACC ATC ATC ACC TTC CTT -3' |
| OMT87go (FP) | *PDF1.3* | 5'- TCT GCT GCC ATC ATC ACT TT -3' |
| OMT88go (RP) | *PDF1.3* | 5'- ACT TCT GTG CTT CCA CCA TTA TCG G -3' |
| OMT93go (FP) | *AT1G67865* | 5'- ATC ATT ACT GGG TGT GGT CAG -3' |
| OMT94go (RP) | *AT1G67865* | 5'- GGA AAC TAC GGC AAT GGT AAT G -3' |
| OMT109go (FP) | *AT1G59780* | 5'- CTG GTT CTT GTG GCC AGT TA -3' |
| OMT110go (RP) | *AT1G59780* | 5'- GAA GGT ATC CAA AGG AGG AGA AG -3' |
| OMT157 (FP) | *ECS1* | 5'- AGT CTC TTC CAT GTT TCT CTT TCT -3' |
| OMT158 (RP) | *ECS1* | 5'- AGT TCA CGG AGT TCC ATT CG -3' |
| OMT163 (FP) | *FMO1* | 5'- CGT CCC ATC TTC AAA CAC AAT C -3' |
| OMT164 (RP) | *FMO1* | 5'- CTT GTC AAA TGG CGA TCA TAC C -3' |
| OMT165 (FP) | *WRKY54* | 5'- CTG TCG AAG AGC CCT CAT AAT AA -3' |
| OMT166 (RP) | *WRKY54* | 5'- CAG CAG TGT GAC AGC AAT AGA -3' |
| OMT175 (FP) | *WRKY70* | 5'- ACG TGT GGT TTC CTA TGT ATG T -3' |
| OMT176 (RP) | *WRKY70* | 5'- AAG TAT ACC CAA GGG TGC AAG -3' |
| OMT268 (FP) | *SMN2* promoter | 5'- TGA TTA CGC CAA GCT TGA CAA TAA AGA TAA ATG GTG AAC TAA G -3' |
| OMT270 (RP) | *SMN2* promoter | 5'- CCG GGG ATC CTC TAG AGG ATT TCT TCA AAA AGA AAA CTT TTA ATC TGT -3' |
