## Supplemental Figures_S1-S5 for "*Arabidopsis SMN2/HEN2*, Encoding DEAD-box RNA Helicase, Governs Proper Expression of the Resistance Gene *SMN1*/*RPS6* and Is Involved in Dwarf, Autoimmune Phenotypes of *mekk1* and *mpk4* Mutants"

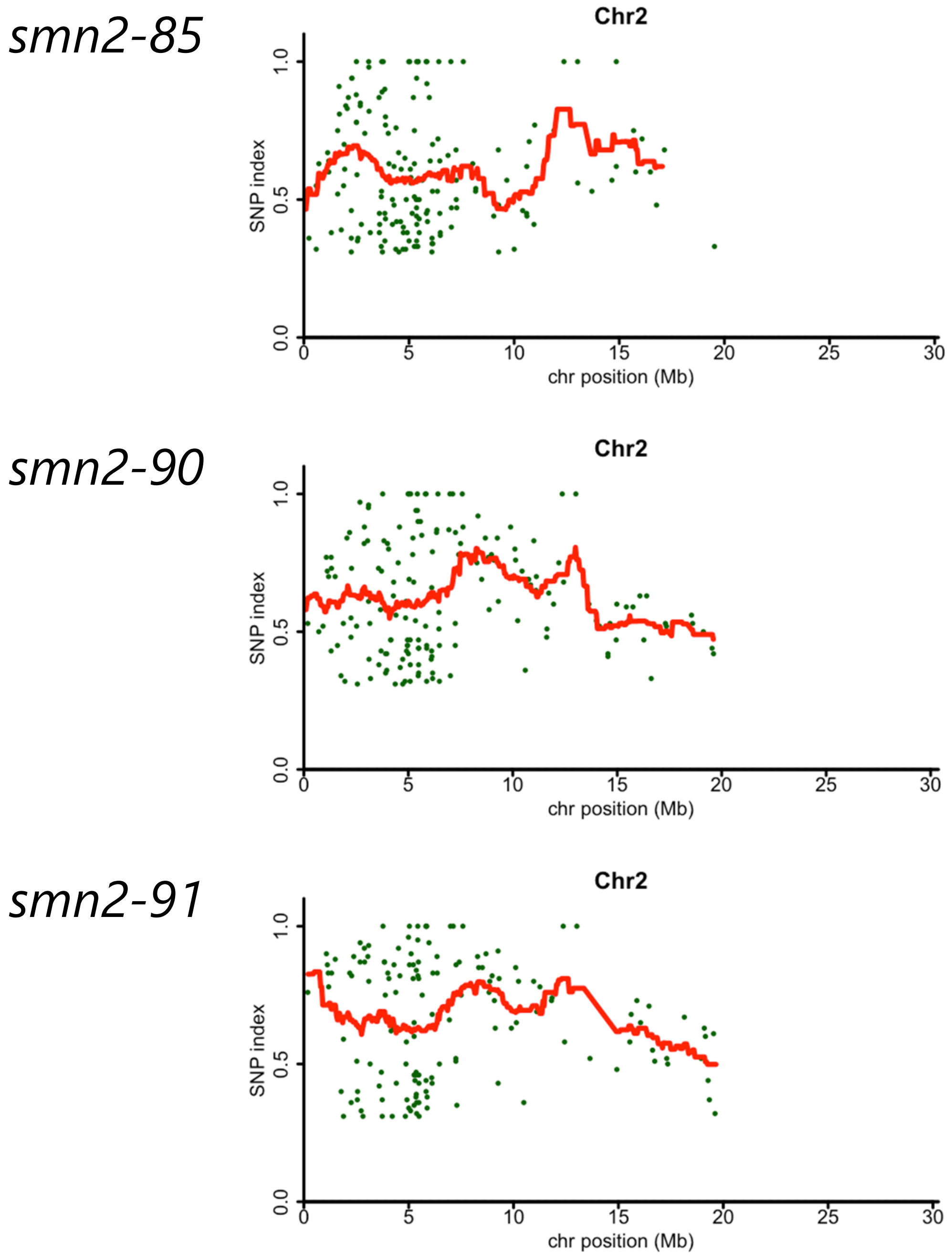


**Supplemental Figure S1. SNP index plots of chromosome 2 from the MutMap analysis of *smn2* mutant alleles.**
Green points, individual SNPs. Red lines, sliding-window average of the ΔSNP index (window size, 2 Mb; slide size, 200 kb).

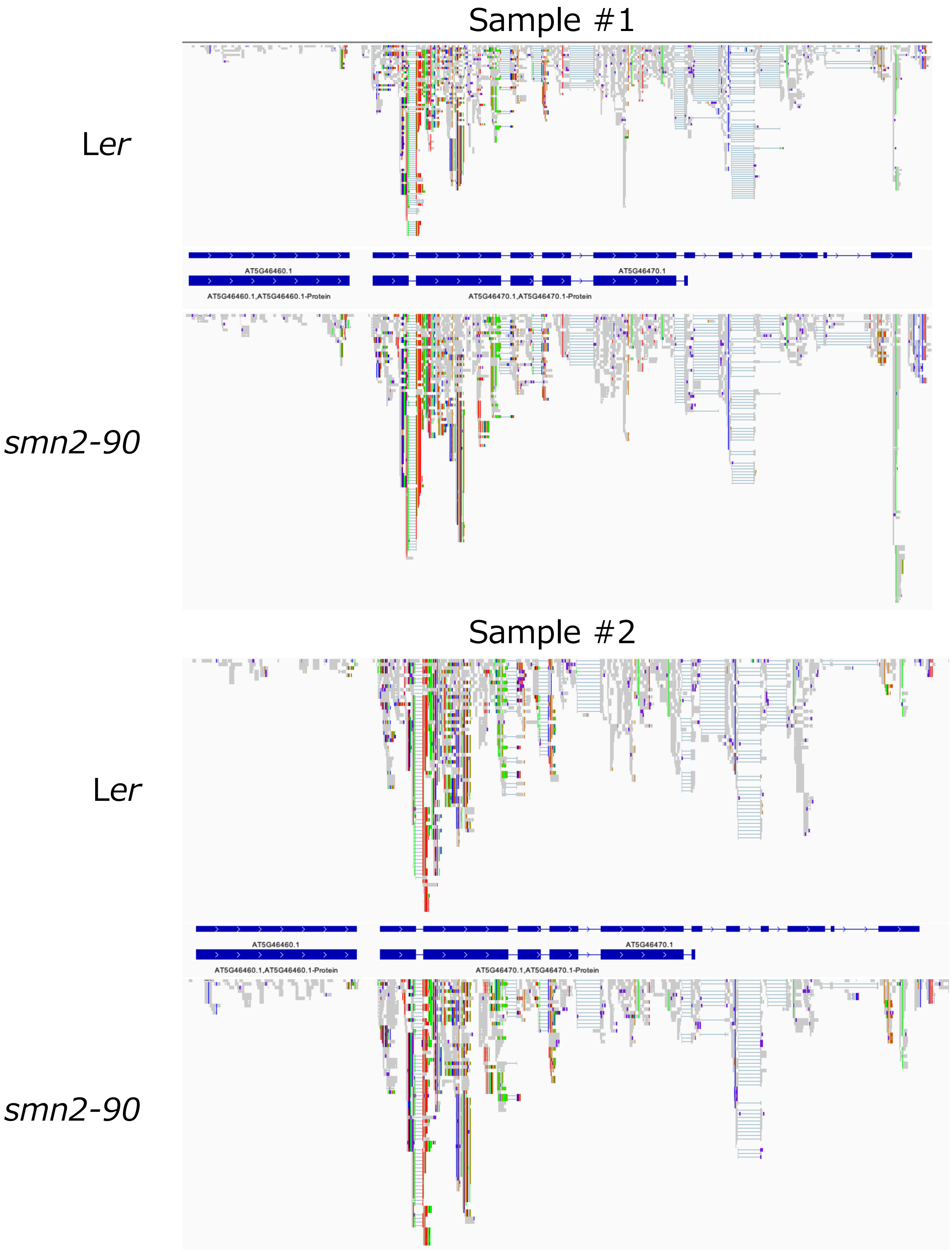


**Supplemental Figure S2. Accumulation of aberrant transcripts of the 3′ region of *SMN1* in *smn2-90* and L*er*.**

Screenshots of RNA-seq data mapped to the *SMN1*/*RPS6* region of the reference *Arabidopsis* genome (TAIR 10) visualized in the Integrative Genomics Viewer (version 2.7.2). Two biological replicates of each L*er* and *smn2-90* were used for RNA-seq. The gene models of *SMN1*/*RPS6* (top) and its coding region (bottom) are indicated in indigo blue, where boxes are exons and lines are introns. Stacked gray objects, reads; stacked sky-blue objects, gaps. Purple dots, deletions; dots of other colors, substitutions vs. reference genome.

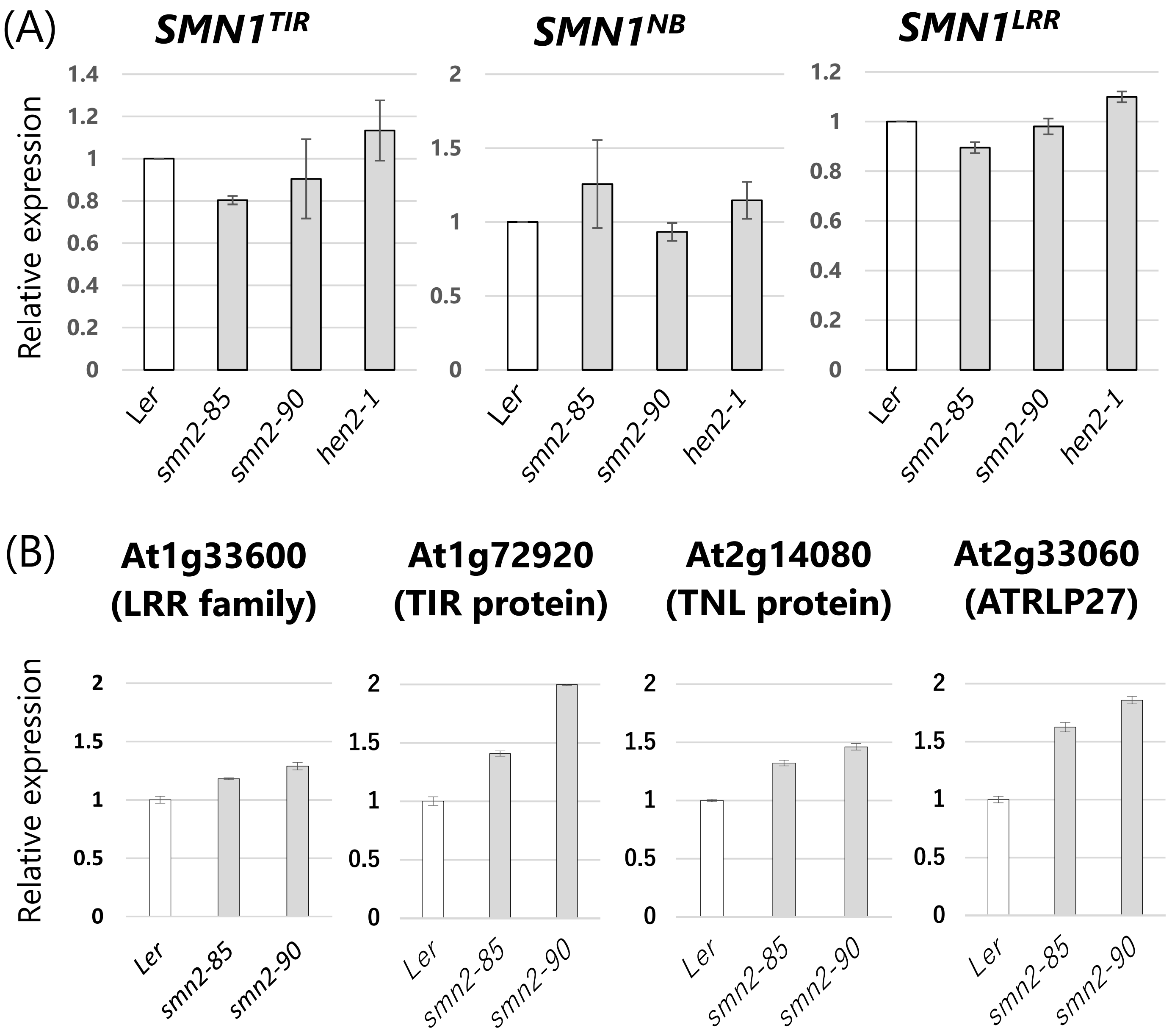


**Supplemental Figure S3. Levels of transcripts of the coding regions of *SMN1/RPS6* and affected defense genes in *smn2* and *hen2-1* mutants.**

(A) Transcripts encoding the TIR, NB, and LRR domains of SMN1/RPS6 (see Fig. 3A). (B) Transcripts of the coding regions of affected defense genes. RNA was extracted from 10- or 14-day-old seedlings, and the transcript levels were analyzed by RT-qPCR. Transcript levels are shown relative to those in L*er*; *Actin2* was used as an internal standard. Error bars, SD (n = 3). A representative result from three independent experiments is shown.

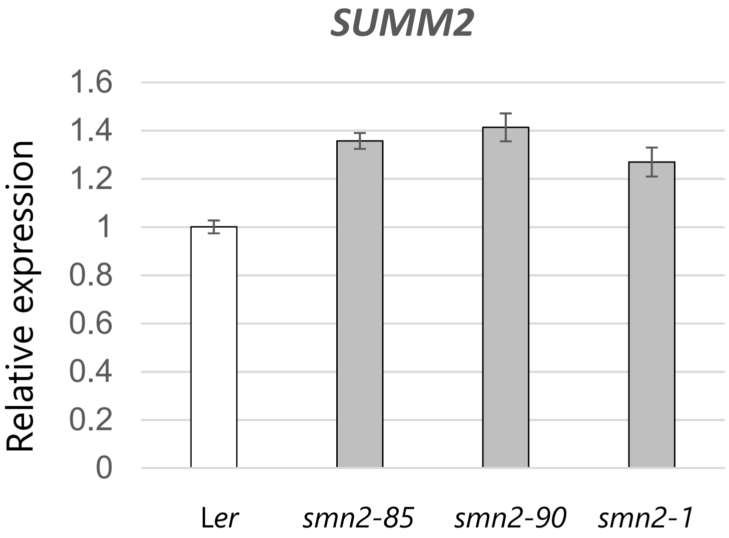


**Supplemental Figure S4.** ***SUMM2*** **transcript levels in** ***smn2*** **and** ***hen2-1*** **mutants.**
Transcript levels of the *SUMM2* coding region near the 3′ end were analyzed by RT-qPCR using RNA extracted from 10-day-old seedlings. Transcript levels are shown relative to those in L*er*; *Actin2* was used as an internal standard. Error bars, SD (n = 3). A representative result from three independent experiments is shown.

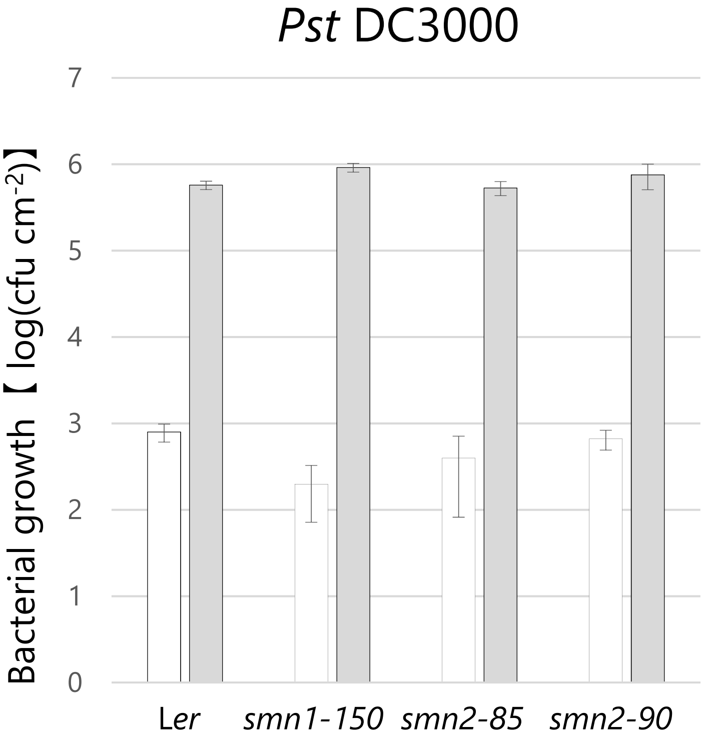


**Supplemental Figure S5. Population growth of *Pst* DC3000 in *smn2* mutants.**
Plants (7 to 8 week old) were inoculated at $1 \times 10^5$ cfu/mL. Values are averages for leaf tissue 0 day (white bar) and 3 days (gray bar) after inoculation. Error bars, SD (n =3). This experiment was performed three times with similar results.
