## Supplementary figures and images for "*Arabidopsis SMN2/HEN2*, Encoding DEAD-box RNA Helicase, Governs Proper Expression of the Resistance Gene *SMN1*/*RPS6* and Is Involved in Dwarf, Autoimmune Phenotypes of *mekk1* and *mpk4* Mutants"

### Supplemental Figure S6

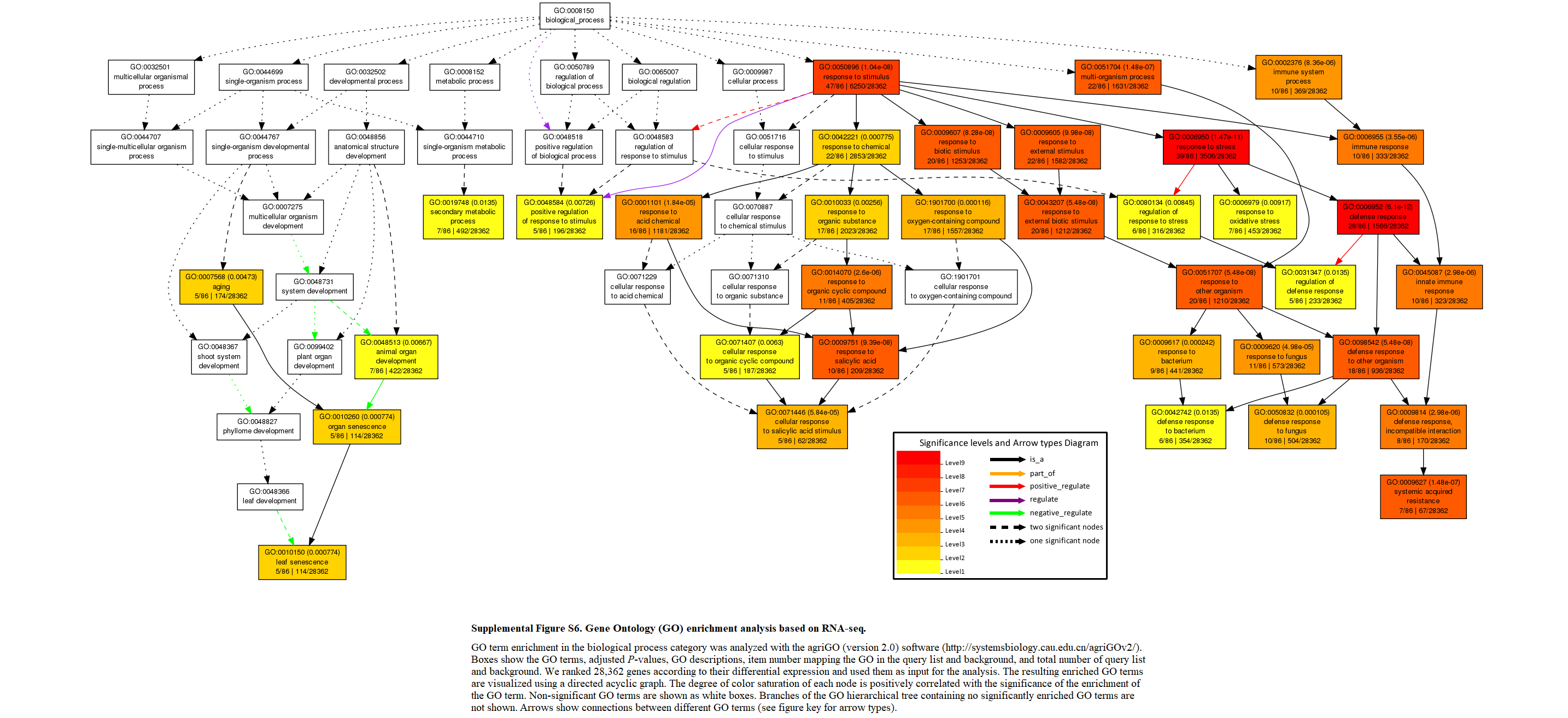
